# Targeting lipid nanoparticles to the blood brain barrier to ameliorate acute ischemic stroke

**DOI:** 10.1101/2023.06.12.544645

**Authors:** Jia Nong, Patrick M. Glassman, Sahily Reyes-Esteves, Helene C. Descamps, Vladimir V. Shuvaev, Raisa Y. Kiseleva, Tyler E. Papp, Mohamad-Gabriel Alameh, Ying K. Tam, Barbara L. Mui, Serena Omo-Lamai, Marco E. Zamora, Tea Shuvaeva, Evguenia Arguiri, Christoph A Thaiss, Jacob W. Myerson, Drew Weissman, Scott E. Kasner, Hamideh Parhiz, Vladimir R. Muzykantov, Jacob S. Brenner, Oscar A. Marcos-Contreras

## Abstract

After more than 100 failed drug trials for acute ischemic stroke (AIS), one of the most commonly cited reasons for the failure has been that drugs achieve very low concentrations in the at-risk penumbra. To address this problem, here we employ nanotechnology to significantly enhance drug concentration in the penumbra’s blood-brain barrier (BBB), whose increased permeability in AIS has long been hypothesized to kill neurons by exposing them to toxic plasma proteins. To devise drug-loaded nanocarriers targeted to the BBB, we conjugated them with antibodies that bind to various cell adhesion molecules on the BBB endothelium. In the transient middle cerebral artery occlusion (tMCAO) mouse model, nanocarriers targeted with VCAM antibodies achieved the highest level of brain delivery, nearly 2 orders of magnitude higher than untargeted ones. VCAM-targeted lipid nanoparticles loaded with either a small molecule drug (dexamethasone) or mRNA (encoding IL-10) reduced cerebral infarct volume by 35% or 73%, respectively, and both significantly lowered mortality rates. In contrast, the drugs delivered without the nanocarriers had no effect on AIS outcomes. Thus, VCAM-targeted lipid nanoparticles represent a new platform for strongly concentrating drugs within the compromised BBB of penumbra, thereby ameliorating AIS.

**Graphical abstract:** 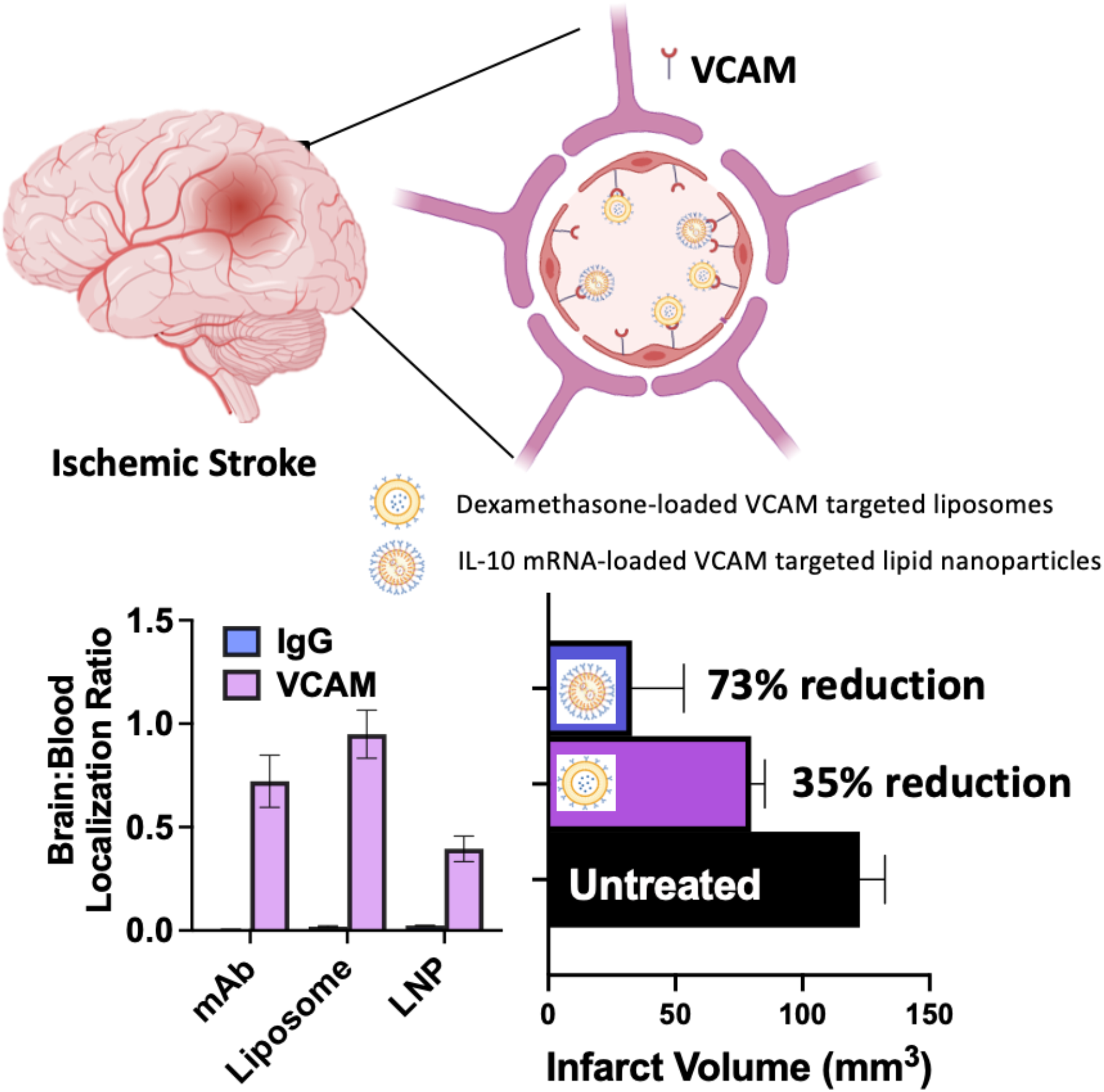

Acute ischemic stroke induces upregulation of VCAM. We specifically targeted upregulated VCAM in the injured region of the brain with drug- or mRNA-loaded targeted nanocarriers. Nanocarriers targeted with VCAM antibodies achieved the highest brain delivery, nearly orders of magnitude higher than untargeted ones. VCAM-targeted nanocarriers loaded with dexamethasone and mRNA encoding IL-10 reduced infarct volume by 35% and 73%, respectively, and improved survival rates.

## Introduction

Stroke is the second leading cause of death and top cause of major disability globally ^1^; however, there have been no new drug classes approved to treat stroke in over 25 years. The vast majority (87%) of strokes are classified as acute ischemic stroke (AIS), characterized by occlusion of an artery supplying the brain ^2^. Treatment of AIS is focused solely on recanalization and reperfusion of the brain (e.g., removal of the clot), which is performed by enzymatic thrombolysis using tissue-type plasminogen activator (tPA) and derivatives or, since 2015, mechanical thrombectomy ^3-6^. The advent of mechanical thrombectomy has provided great improvement in restoring blood flow to ischemic regions of the brain; however, outcomes of severe AIS (characterized by large vessel occlusion) are poor, with 50-55% of patients dying or becoming functionally dependent on others ^2,3,6^. In order to improve outcomes, decades of research has been performed with the goal of salvaging the reperfused or partially perfused (penumbra) tissue surrounding the infarct, namely that caused by ischemia-reperfusion injury. A diverse array of drugs have been tested that target specific mechanisms of this injury (e.g., excitotoxicity, neuronal apoptosis, neuroinflammation), with particular promise being shown for inhibition of neuroinflammation in preclinical studies ^7^. Neuroinflammatory processes are propagated by expression of intercellular adhesion molecule 1 (ICAM) and vascular cell adhesion molecule 1 (VCAM) by endothelial cells in the penumbra, which help recruit immune cells to the injured region ^8-11^.

Despite promising results in preclinical studies, all neuroprotectants that progressed to clinical trials have failed in development, with the primary cause of failure being cited as insufficient drug delivery ^12^. Several features of AIS make it a particular challenge for drug delivery. First, efficient drug delivery to the brain is notoriously difficult in general due to the nearly impenetrable blood-brain barrier. Second, poor reperfusion (common prior to mechanical thrombectomy) leads to insufficient blood flow to at-risk brain regions to achieve effective drug accumulation. Third, severe AIS patients are critically ill and therefore do not tolerate off-target drug effects, resulting in strict limitations on systemic dosing in AIS. Therefore, there is a critical unmet need to improve drug delivery to the brain in AIS, which we hypothesize can be achieved using targeted drug delivery systems.

Nanocarriers (NCs) are particles with diameters of ∼100 nm that allow the encapsulation and delivery of diverse small molecule drugs and biotherapeutics. Clinical use of NCs originated in the field of oncology with the approval of liposomal doxorubicin (Doxil, Caelyx) in the 1990s. NCs have recently risen to prominence in the form of COVID-19 vaccines (Comirnaty, Spikevax), which used lipid nanoparticles (LNP) to encapsulate mRNA, with demonstrated safety and efficacy in a global population (>5 billion doses administered) ^13^. Selective delivery of NCs to tissues, sites of injury, or specific cells can be achieved by conjugation of affinity ligands on the NC surface. This targeted nanomedicine allows: 1) localization and concentration of drugs in the target site; and, 2) reduction in off-target toxicities by sparing tissues from drug exposure, thereby increasing the therapeutic index ^14^. We hypothesized that targeted, clinically relevant NCs would be useful for the treatment of AIS.

It is well-established that inducible cell-surface molecules such as ICAM and VCAM are upregulated in the penumbra in AIS as a result of neuroinflammation ^15-17^. Therefore, we hypothesized that targeting these inducible markers would concentrate our NCs in the injured region of the brain, and provide therapeutic efficacy that free drugs could not achieve. In the current study, we used the gold standard animal model of large vessel AIS followed by mechanical thrombectomy (transient middle cerebral artery occlusion, tMCAO). This model is characterized by occlusion of the middle cerebral artery (MCA) with a filament for a pre-defined period of time, followed by filament removal and reperfusion. We have demonstrated that, in this model: 1) comparison of ICAM and VCAM accessibility to IV-injected monoclonal antibodies (mAb) and NCs (liposomes and mRNA-containing LNP), as well as the ones targeted to constitutively expressed platelet endothelial CAM (PECAM); 2) direct quantification of radiolabeled mAbs and NCs that showed a superior targeting specificity to injured brain for VCAM-directed agents; 3) uptake of VCAM-targeted NCs in endothelial cells and infiltrating leukocytes; 4) expression of reporter cargoes (luciferase) and gene editing machinery (Cre-recombinase) in the injured region of the brain following IV dosing of VCAM-targeted LNP; 5) a 35% reduction in infarct volume following IV dosing of VCAM-targeted, dexamethasone-loaded liposomes; a 73% reduction of infarct volume with a 100% survival following intravenous administration of mRNA LNP encoding of very potent anti-inflammatory cytokine Interleukin 10 (IL-10). In sum, these results demonstrated that VCAM targeting is a useful approach to improve the efficacy of neuroprotective drugs in a mouse model of AIS.

## Materials and Methods

### Reagents

All lipids for liposome preparation were purchased from Avanti Polar Lipids (Alabaster, AL). LNP were provided by Acuitas Therapeutics (Vancouver, Canada). Dibenzocyclooctyne (DBCO)-PEG4-NHS ester was purchased from Click Chemistry Tools (Scottsdale, AZ). Na^125^I was purchased from PerkinElmer (Waltham, MA). Rat IgG and antibodies for flow cytometry were purchased from Invitrogen (Carlsbad, CA). Anti-mouse-PECAM-1 (clone MEC13.3) was purchased from BioLegend (San Diego, CA). All other chemicals and reagents were purchased from SigmaAldrich (St. Louis, MO), unless otherwise noted.

### Animals

All animal studies were carried out in accordance with the *Guide for Care and Use of Laboratory Animals* under the protocol approved by the University of Pennsylvania Institutional Animal Care and Use Committee. All animal experiments were carried out using male, 6-8 week-old C57BL/6 mice (20-25g) from Jackson Laboratory (Bar Harbor, ME), unless otherwise noted. Ischemic stroke was induced using the tMCAO model [18]. Mice were randomly assigned to groups and anesthetized with isoflurane. After proper anesthetic depth was verified, the mouse was placed on a heating pad with rectal temperature maintained at 37°C. The carotid artery was exposed under a dissecting microscope. A small arteriotomy was made and a filament was threaded into the MCA. A doppler probe was put over the MCA. Greater than 70% reduction of MCA blood flow was confirmed by Doppler ultrasound as the criteria for successful MCAO. Forty-five minutes later, the filament was removed, MCA blow flow return was confirmed by Doppler, and hemostasis was achieved. Greater than 80% of blood flow restoration within 10 minutes after filament removal was required for animal inclusion in the study. The mouse was returned to its cage, and given 1 mL of normal saline subcutaneously to maintain hydration daily. Subsequently, food and water were provided on the floor of cage in a petri dish.

### Protein Production and Purification

Anti-mouse-VCAM-1 mAb (clone M/K2.7) and anti-mouse-ICAM-1 mAb (clone YN1/1.7.4) were produced by culturing hybridoma cells and purified using protein G sepharose chromatography (GE Healthcare Life Sciences, Pittsburgh, PA).

### Antibody Radiolabeling

Antibodies were radiolabeled with Na^125^I using Pierce Iodogen radiolabeling method. Briefly, tubes were coated with 100 μg of Iodogen reagent. Antibody (1-2 mg/mL) and Na^125^I (0.25 μCi/μg protein) were incubated for 5 minutes on ice. Unreacted materials were purified using Zeba 7 kDa desalting spin columns (ThermoFisher Scientific). Thin layer chromatography was used to confirm radiolabeling efficiency. All proteins were confirmed to have >90% radiochemical purity prior to use.

### Antibody Modification

Antibodies were functionalized with DBCO or N-succinimidyl S-acetylthioacetate (SATA) for conjugation to liposomes or LNP, respectively, by reacting with a 5-fold molar excess of DBCO-PEG4-NHS ester or SATA for 30 minutes at room temperature. SATA deprotection was carried out with 0.5 M hydroxylamine. The unreacted compound was removed via centrifugation using a molecular weight cutoff filter or G-25 Sephadex Quick Spin Protein columns (Roche Applied Science, Indianapolis, IN).

### Liposome Preparation

Azide functionalized liposomes were prepared by thin film hydration techniques, as previously described [19]. Briefly, 1,2-dipalmitoyl-sn-glycero-3-phosphocholine (DPPC), cholesterol, and 1,2-distearoyl-sn-glycero-3-phosphoethanolamine-N-[azido(polyethyleneglycol)-2000 (DSPE-PEG2000-azide) were mixed in a molar ratio of 54:40:6. For flow cytometry studies, fluorescent liposomes were prepared by doping in a 0.1% molar ratio of TopFluor PC into the lipid film. The lipid film was rehydrated in phosphate-buffered saline (PBS), pH 7.4. To form drug-loaded liposomes, the lipid film was hydrated in a solution containing 40 mg/mL of dexamethasone-21-phosphate (Dex) in PBS, pH 7.4. The resulting vesicles were extruded through 200 nm polycarbonate membranes. The drug entrapment efficiency was assessed using reversed phase high performance liquid chromatography (HPLC), using a mobile phase containing 30% v/v acetonitrile, 70% v/v water, and 0.1% v/v trifluoroacetic acid and running through a C8 column (Eclipse XDB-C8, 3 μm, 3.0×100 mm, Phenomenex) at a flow rate of 0.6 mL/minute. Dex was detected using UV absorbance at 240 nm.

### Preparation of mRNA-LNP

mRNAs were produced as described previously [20] using T7 RNA polymerase (Megascript, Ambion) on linearized plasmids encoding codon-optimized NanoLuc Luciferase (Promega), Cre recombinase, or murine IL-10 (NCBI Reference Sequence: NP_034678.1). To make modified nucleoside-containing mRNA, m1Ψ-5′-triphosphate (TriLink) was incorporated instead of UTP. mRNAs were transcribed to contain 101 nucleotide-long poly(A) tails. They were capped using the m7G capping kit with 2′-O-methyltransferase (ScriptCap, CellScript) to obtain cap1. mRNA was purified by Fast Protein Liquid Chromatography (FPLC) (Akta Purifier, GE Healthcare) [21]. All prepared RNAs were analyzed by electrophoresis using denaturing or native agarose gels and stored at -20°C. FPLC-purified m1Ψ-containing NanoLuc luciferase and Cre-encoding mRNAs were encapsulated in LNP using a self-assembly process as previously described in which an aqueous solution of mRNA at acidic pH is rapidly mixed with a solution of lipids dissolved in ethanol ^18^. The lipids and LNP composition are described in US patent US10,221,127. mRNA-LNP formulations were stored at -80°C at a concentration of mRNA of ∼1 mg/ml. LNP were then modified with DSPE-PEG-maleimide via a post-insertion technique [25]. Briefly, maleimide-functionalized micelles were prepared by mixing DSPE-PEG(2000) and DSPE-PEG-maleimide at a molar ratio of 4:1. Solvent was evaporated and the lipids were rehydrated in PBS at 65°C with vortex. Size of the micelles was 14-20 nm measured by dynamic light scattering. The desired amount of micelles were then mixed with 1mg/mL of LNP and incubated for 3 hours at 37°C to obtain maleimide-functionalized LNP for antibody conjugation.

### Antibody Conjugation

To conjugate targeting ligands, azide-functionalized liposomes were incubated with DBCO-modified antibodies at 4 °C overnight with rotation. For targeted LNP, SATA-antibodies were conjugated to LNP particles via SATA-maleimide conjugation chemistry for 30 minutes at room temperature. Cystine was added to stop the SATA-maleimide reaction. For experiments involving radiotracing, ^125^I-labeled untargeted rat IgG was spiked in at 10% of the total antibody mass added. Immunoliposomes and immunoLNP were purified using gel filtration chromatography to remove unbound antibodies. The size, distribution, and concentration of the NCs were determined using dynamic light scattering using Zetasizer Nano ZS (Malvern Instruments Ltd, Malvern, UK). and nanoparticle tracking analysis using a Nanosight NS300 (Malvern Panalytical, Westborough, MA). The conjugation efficiency for targeted liposomes is ∼85%, and for targeted LNP ∼95%, resulting in ∼50 mAb/NC.

### Biodistribution

For radiotracing experiments, mice were injected intravenously with radiolabeled antibody (5 µg, ∼0.2 mg/kg), immunoliposomes (10 mg/kg lipid, ∼6E11 liposomes/animal), LNP (8µg of mRNA, ∼0.32 mg/kg) 24 hours post-tMCAO reperfusion. Thirty minutes after injection, animals were sacrificed and perfused with 20 mL of ice-cold PBS. Blood was collected in EDTA-coated tubes. Tissue distribution of injected materials was determined by measuring the radioactivity in the blood and other organs using a Wizard 2470 gamma counter (PerkinElmer, Waltham, MA). Tissue uptake was presented as percent injected dose normalized to the mass of tissue (%ID/g tissue).

### Luciferase Transfection

Transgene expression was assessed in mice following injection of LNP-NanoLuc mRNA and tissues were harvested 5 h post-injection and frozen at -80°C. Tissues were homogenized with 1 ml of Luciferase Cell Culture Lysis (Promega) containing protease inhibitor cocktail using PowerLyser 24 (Qiagen) and mixed gently at 4 °C for one hour. The homogenates were then subjected to cycles of freeze/thaw in dry ice/37 °C. The resulting cell lysate was centrifuged for 10 min at 16,000 g at 4 °C and supernatants were separated. Protein concentration was measured by Lowry assay. Luciferase activity was measured in supernatant using Nano-Glo Dual-Luciferase Reporter Assay System (Promega). Luminescence was assayed using a Victor^3^ 1420 Multilabel Plate Counter (Perkin Elmer, Wellesley, MA). The data were presented as luminescence units normalized by mg of protein.

### Flow Cytometry

To obtain single cell suspensions, brains were disaggregated as previously described [26]. Briefly, tissue was enzymatically digested with dispase at 37 °C for 1 hour, followed by addition of 600 U/mL DNase Grade II. Tissue digests were demyelinated using a Percoll gradient. Residual red blood cells were lysed by ACK buffer (Quality Biological, Gaithersburg, MD). Cells were then stained to determine the cellular distribution of VCAM-targeted liposomes in leukocytes (CD31^-^/CD45^+^) vs. endothelial cells (CD31^+^/CD45^-^) (**Supplemental Table 5**). Flow cytometry was performed using an Accuri C6plus (Benton Dickinson, San Jose, CA).

### Histology

mTmG reporter mice B6.129(Cg)-Gt(ROSA)26Sor^tm4(ACTB-tdTomato,-EGFP)Luo^/J on a C57BL/6J background with two-color fluorescent Cre-reporter allele (Jackson Laboratory, Farmington, CT) were used to determine the functional activity of mRNA/LNP. tMCAO animals were IV injected daily with VCAM targeted Cre-recombinase encoding mRNA-loaded LNP for 3 days starting just after reperfusion. At the end of the study, the brain was harvested, frozen in O.C.T compound (Fisher HealthCare) and cryosectioned using cryostat Leica CM 1950 (Leica Biosystems, Nussloch, Germany). Tissue specimen was then fixed with paraformaldehyde, counterstained with DAPI (Molecular Probes, ThermoFisher), and imaged with Leica DM6000 Widefield microscope.

### Therapeutic Studies

To study the therapeutic effects, VCAM-targeted Dex-loaded liposomes or IL-10 mRNA loaded LNP were dosed IV every 24 hours starting just after removal of the filament for total of 3 doses prior to sacrifice, and compared to their controls. Animals were perfused with ice-cold PBS and the brains were removed and sectioned into 1 mm thickness using a rotary hand microtone. The ischemic infarct was evaluated by immunohistochemical analysis using 1% tetrazolium chloride (TTC) staining. The area of non-stained infarct in each slice was measured using ImageJ in a blinded manner. The ischemic infarct volume area was calculated and multiplied by slice thickness and summed.

### Cytokine Measurement

Plasma and tissue were collected after IL-10 mRNA containing LNP treatment. Tissue was homogenized using protease buffer (5mM EDTA, 10mM Tris, and 1:100 dilution of protease inhibitor cocktail (Sigma) in PBS) and lysis buffer (10% Triton X-100 in PBS). After centrifuge, the supernatant was collected and protein level was measured by Lowry assay. The LEGENDplex Mouse Inflammation Panel 13-plex (Biolegend) was used for cytokine quantification. Measurement was performed on an LSR Fortessa B flow cytometer (Becton Dickinson, Franklin Lakes, NJ) using LEGENDplex software for data analysis.

### Statistics

All results are expressed as mean ± SEM. Statistical analyses were performed using GraphPad Prism 8 (GraphPad Software, San Diego, CA). Unless specified, we used 1-way ANOVA for multiple comparisons. * denotes p<0.05, ** denotes p<0.01, *** denotes p<0.001, **** denotes p<0.0001.

## Results

### Targeting the tMCAO brain using monoclonal antibodies directed against CAMs

Both AIS in humans and tMCAO in mice (**Figure 1A**) lead to severe acute inflammation in the penumbra around the core infarct, resulting in upregulation of inducible CAMs (e.g., ICAM and VCAM) ^19,20^. Prior studies have shown this increase in CAM expression in tMCAO by measuring mRNA and protein levels in tissue homogenates, or with immunohistology ^21-23^. These studies left an open question: Are the *de novo* expressed ICAM and VCAM proteins accessible to the vessel lumen, allowing adhesion of targeted therapeutics? To assess vascular accessibility of CAMs and to establish their use for targeted drug delivery in tMCAO and AIS, we injected radiolabeled mAbs against PECAM (constitutively expressed), and ICAM and VCAM (inducible) 24 hours after reperfusion ^24^. Mice were sacrificed and perfused 30-minutes post-injection to determine the tissue distribution of the mAbs (**Figure 1B**).

**Figure 1.**
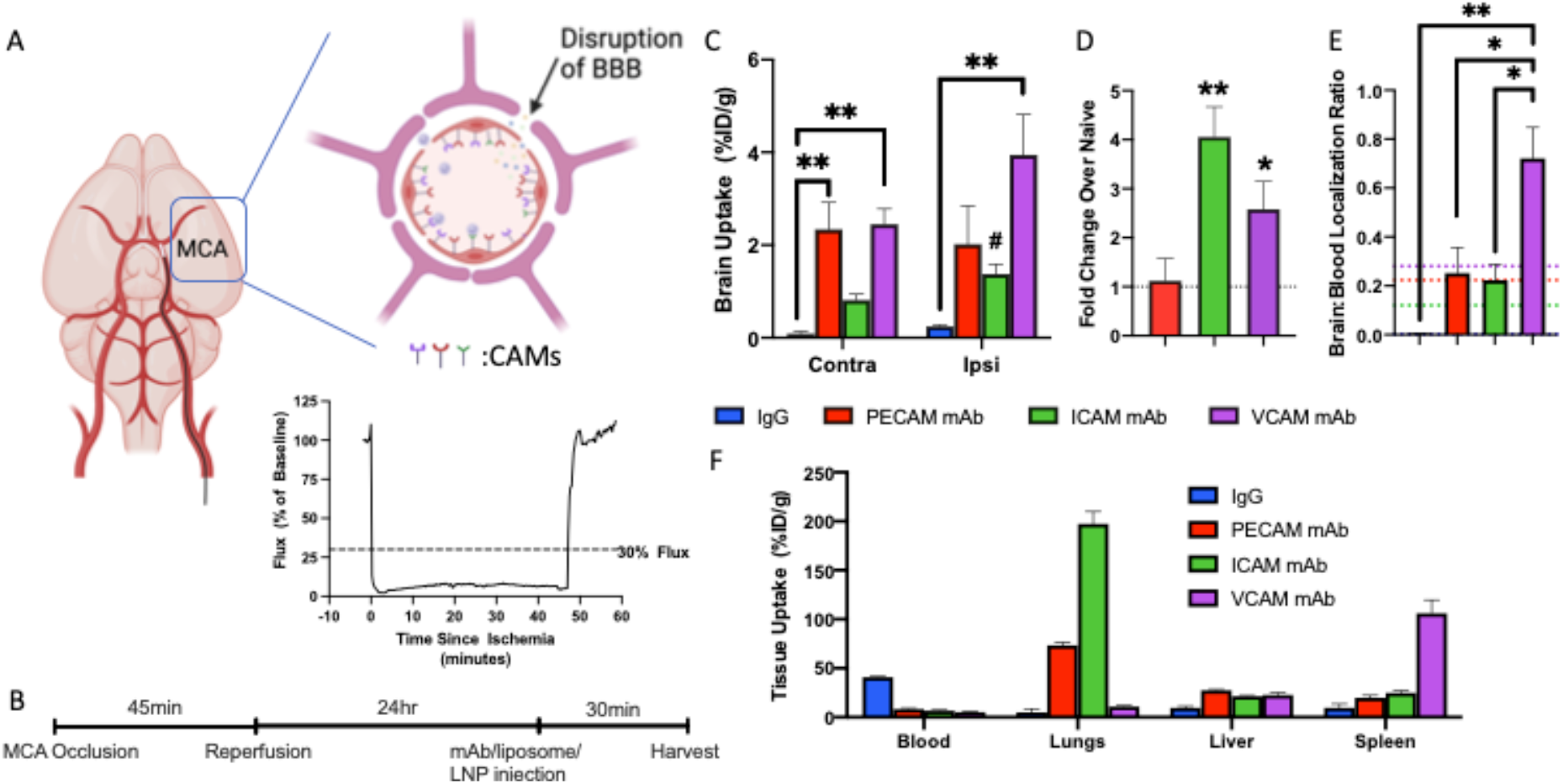
In the tMCAO model of AIS, IV-injected antibodies accumulate in the brain at vastly higher levels when they target cell adhesion molecules (CAMs). (A) Schematic representation of the tMCAO model, where a filament is inserted through the internal carotid until it blocks the middle cerebral artery (MCA). This ischemia induces neuronal cell death in the core infarct; in the penumbra of partially perfused tissue, there is disruption of the BBB and upregulation of luminal expression of CAMs (top right insert). We used a laser Doppler flowmeter to monitor blood flow before and during the ischemic period (t = 0 to 45 minutes), and after reperfusion (bottom right inset, representative animal). More than 80% blood flow restoration within 10 minutes after filament removal was taken as criteria for reperfusion, required for animal inclusion in the study. (B) Timeline for all biodistribution experiments. Mice were challenged with ischemia (45 min) and reperfusion, followed by IV injection of ^125^I-radiolabeled mAbs/liposomes/LNP or untargeted IgG controls 24 hours post-injury, and tissue biodistribution was determined 30 min post-injection. (C) All anti-CAM mAbs had markedly higher brain uptake than control IgG, with the highest uptake in the ipsilateral (injured) hemisphere being anti-VCAM. Anti-ICAM mAb was the only mAb that showed a statistically significant preference for accumulation in the ipsilateral vs. contralateral hemisphere. (D) Fold change over naïve in ipsilateral tMCAO brain showed upregulation of ICAM and VCAM after injury. (E) The brain-to-blood distribution ratio in ipsilateral tMCAO brain further illustrates the superiority of VCAM-targeted mAbs. The values observed in naïve mice are shown by dashed lines. (F) Biodistributions to blood, lungs, liver, and spleen are shown for the mAbs in tMCAO mice. Note the high lung uptake for mAbs targeted to ICAM and PECAM, while anti-VCAM was mainly concentrated in the spleen, and untargeted IgG control remains in circulation (high blood level). N=3. *p<0.05, **p<0.01 using one-way ANOVA followed by Dunnett’s post-hoc against IgG, #p<0.05 using paired t-test comparing contralateral vs. ipsilateral.

In tMCAO mice, CAM-targeted mAbs accumulated in the brain to a significantly higher degree than control IgG (**Figure 1C**), precluding vascular leak as a driver of tissue uptake. Anti-VCAM had the highest brain uptake at 3.95 ± 0.87 %ID/g in the ipsilateral (injured) hemisphere, while IgG control had the lowest uptake (0.25 ± 0.03 %ID/g). There was no significant difference between the two hemispheres for anti-PECAM (2.34 ± 0.05 %ID/g contralateral vs. 2.02 ± 0.82 %ID/g ipsilateral) and anti-VCAM mAb (2.45 ± 0.33 %ID/g contralateral vs. 3.95 ± 0.87 %ID/g ipsilateral), but anti-ICAM had a small but statistically significant preference for the ipsilateral hemisphere (0.82 ± 0.12 %ID/g contralateral vs. 1.38 ± 0.21 %ID/g ipsilateral) (**Figure 1C**). The fold change of CAM expression in ipsilateral tMCAO brain over naïve showed significant upregulation of ICAM and VCAM (**Figure 1D**). To better understand the uptake of the brain, a more relevant term “localization ratio” or “blood-normalized tissue uptake” is used to determine the brain-to-blood ratio. This helps to demonstrate whether the accumulation is caused by high blood concentrations leaking into the brain, which also produce continued side effects by leaking cargo drug to other tissues. As shown in **Figure 1E**, in ipsilateral brain of tMCAO mice, the blood-normalized brain uptake of anti-VCAM mAb was more than two-fold higher (0.72 ± 0.13) than anti-PECAM (0.25 ± 0.10) and anti-ICAM mAb (0.22 ± 0.07) and exceeded IgG uptake by more than two orders of magnitude (0.01 ± 0.001). Additionally, compared to naïve mice (no tMCAO), only the brain-to-blood uptake of VCAM increased after tMCAO challenge. VCAM mAbs outperformed mAbs for other CAMs in magnitude of brain uptake and in augmentation of uptake by AIS pathology.

We also investigated CAM-targeted mAbs uptake in other organs (**Figure 1F**). This analysis showed that anti-PECAM and anti-ICAM mAbs primarily accumulated in the lungs (73.4 ± 2.9 % ID/g and 197.4 ± 12.4 % ID/g, respectively), anti-VCAM mAb accumulated in the spleen (106.2 ± 13.5 % ID/g), and untargeted IgG maintained high concentration in the blood (41.0 ± 0.82 % ID/g). This is consistent with observations in previous publications with acute brain inflammation (**Supplemental Table 1**) ^25^.

**Table 1.** Characterization of targeted NCs. N=3. All values represented as mean ± SEM of three independent measurements.

| Size | IgG | PECAM | ICAM | VCAM |
| --- | --- | --- | --- | --- |
| LNP | 89 ± 2 | 92 ± 2 | 86 ± 2 | 89 ± 0.3 |
| Liposomes | 153 ± 3 | 155 ± 2 | 159 ± 5 | 159 ± 3 |
| PDI |  |  |  |  |
| LNP | 0.055 ± 0.006 | 0.067 ± 0.011 | 0.052 ± 0.007 | 0.047 ± 0.012 |
| Liposomes | 0.112 ± 0.023 | 0.131 ± 0.011 | 0.116 ± 0.008 | 0.12 ± 0.08 |

### VCAM targeting outperforms ICAM and PECAM targeting for mRNA-LNP and liposome delivery to tMCAO brains

Having shown that CAM targeting directs mAbs to the tMCAO brain, we examined CAM targeting with mRNA-loaded LNP and liposomes. Compared to mAbs, NCs have vastly different accessibility to membrane epitopes (due to steric hindrance), pharmacokinetics, and binding capacity. We characterized two NCs that are FDA approved and widely used in clinical studies: LNP and liposomes. Liposomes have potential for delivering a wide array of small molecule and protein drugs, including dexamethasone ^12,26^ and LNP allow delivery of RNA cargoes (e.g., mRNA or siRNA).

We evaluated brain uptake of LNP directed against PECAM, ICAM, and VCAM, compared to untargeted IgG control (**Figure 2A and Table 1**). Similar to CAM-targeted mAbs, within 30 minutes of injection, both PECAM- and VCAM-targeted LNP accumulated in the brain at significantly higher concentrations than untargeted IgG LNP. Ipsilateral to contralateral ratios showed a higher selectivity for the ipsilateral part of ICAM and VCAM targeted NCs compared to PECAM (ipsilateral/contralateral ratios: ICAM 1.91 ± 0.43, VCAM 1.54 ± 0.14 and PECAM 1.22 ± 0.08) (**Figure 2B**). When comparing localization ratio of all LNP in the ipsilateral brain, uptake of anti-VCAM LNP achieved 16.5-fold superiority over untargeted IgG LNP (0.40 ± 0.1 vs. 0.02 ± 0.002), 2.23-fold superiority over anti-PECAM LNP (0.18 ± 0.01), and 3.91-fold superiority over anti-ICAM LNP (0.10 ± 0.01) (**Figure 2C**). As expected, anti-PECAM and anti-ICAM LNP also accumulated in the lungs and the absolute delivery to the pulmonary vasculature of these formulations greatly exceeded that in the brain (pulmonary uptake of anti-PECAM and anti-ICAM LNP achieved 85.0 ± 2.2 % ID/g, 172.7 ± 21 % ID/g, respectively). Anti-VCAM LNP minimally accumulated in the lungs, and were largely taken up in the spleen (113.8 ± 7 % ID/g, **Figure 2D and Supplemental Table 2**).

**Figure 2.**
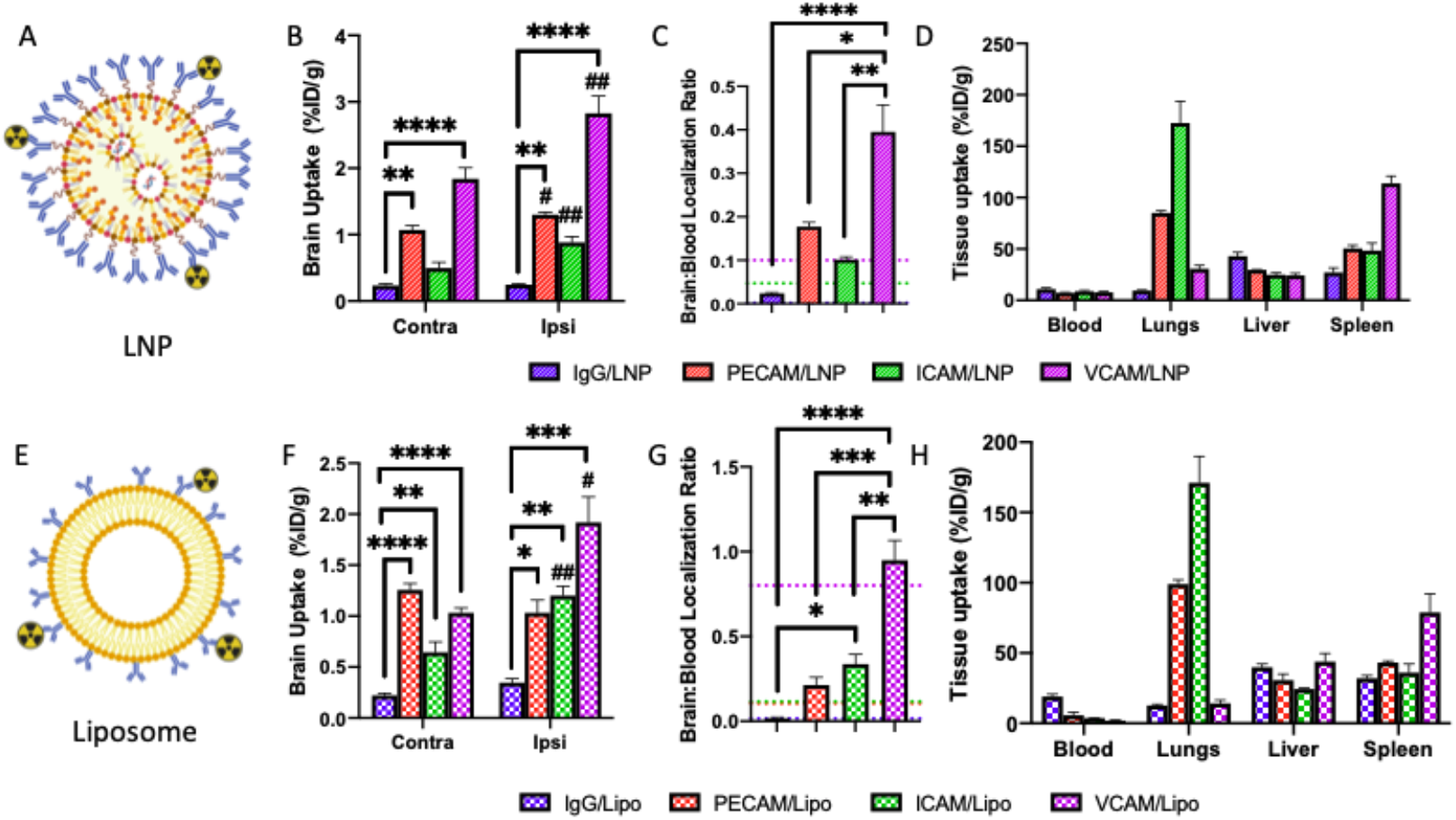
CAM-targeting increases NC accumulation in the inflamed brain after tMCAO ischemia-reperfusion injury in the order VCAM>ICAM=PECAM>>IgG. (A) Schematic representation of radiolabeled, antibody-decorated LNP-mRNA. (B) Brain biodistributions showed specific accumulation of CAM-targeted LNP in the post-tMCAO brain, especially in the ipsilateral hemisphere. (C) VCAM targeting significantly enhanced blood-normalized ipsilateral brain uptake of LNP, compared to other tested CAMs or untargeted control. (D) LNP biodistribution to major organs 30 minutes after injection: clearance from the blood of all CAM-targeted LNP and untargeted IgG control LNP; ICAM- and PECAM-targeted LNP accumulated at highest concentration in the lungs, while VCAM-targeted LNP were mainly taken up by the spleen. (E) Schematic representation of radiolabeled immunoliposomes. (F) Brain uptake of anti-CAM liposomes in post-tMCAO mice was markedly higher than untargeted (control) IgG. Both ICAM- and VCAM-targeted liposomes had significantly higher accumulation in the ipsilateral hemisphere compared to the contralateral one. (G) VCAM-targeted liposomes had higher blood-normalized ipsilateral brain uptake compared to other CAM-targeted liposomes and IgG control. (H) As with targeted LNP, targeted liposomes rapidly cleared from the bloodstream, ICAM and PECAM targeting achieved high lung uptake, and VCAM targeting achieved high splenic uptake. N=3-6. Dashed lines indicate localization ratio of targeted NC in naïve brains. *p<0.05, **p<0.01, ***p<0.001, ****p<0.0001 using one-way ANOVA followed by Dunnett’s posthoc test, #p<0.05 and ##p<0.01 using paired t-test contralateral vs. ipsilateral.

The CAM-targeted LNP study design was replicated to assess brain uptake of CAM-targeted immunoliposomes (**Figure 2E and Table 1**). CAM targeting significantly enhanced liposome uptake in both hemispheres of tMCAO-challenged brains, compared to untargeted IgG control (**Figure 2F)**. ICAM- and VCAM-targeted liposomes had preferential delivery to the ipsilateral hemisphere in tMCAO (ICAM: 0.6 ± 0.1 %ID/g contralateral vs. 1.2 ± 0.1 %ID/g ipsilateral, VCAM: 1.0 ± 0.05 %ID/g contralateral vs. 1.9 ± 0.2 %ID/g ipsilateral). When comparing localization ratio, VCAM-targeted liposomes had 50-fold higher ipsilateral accumulation vs. IgG control (0.95 ± 0.12 vs. 0.02 ± 0.004), 4.46-fold higher than PECAM-targeted liposomes (0.21 ± 0.05), and 2.82-fold higher than ICAM-targeted ones (0.34 ± 0.06) (**Figure 2G**). We observed high pulmonary uptake of anti-PECAM and anti-ICAM liposomes (99.1 ± 3 % ID/g and 171.2 ± 18.8 % ID/g, respectively) and high splenic uptake of anti-VCAM liposomes (78.9 ± 13.1 % ID/g) (**Figure 2H, Supplemental Table 3**). Therefore, anti-CAM targeted delivery systems tested in this study demonstrate reproducible patterns of the uptake in the main organs: PECAM targeting aims to the lungs similarly in healthy and stroke mice; B) ICAM counterparts accumulates preferably to inflammation altered sites, and C) VCAM targeting eludes the lungs and allows these NCs accumulate locally in the inflamed BBB.

These studies show that targeting to PECAM, ICAM, or VCAM can enhance tMCAO brain uptake of LNP and liposomes. Specifically, VCAM targeting produces the best tMCAO brain delivery by nearly every metric, exceeding untargeted NCs by 1-2 orders of magnitude. Therefore, additional experiments in this study are focused on VCAM targeting.

### VCAM-targeted NCs deliver to the endothelial cells and leukocytes of the tMCAO brain

Having shown strong brain uptake of VCAM-targeted NCs, we determined which cell types in the brain were responsible for this delivery. We injected fluorophore-labeled, VCAM-targeted liposomes intravenously into tMCAO mice, then harvested and disaggregated the brain to create a single cell suspension, following the experimental timeline in **Figure 1B**. Cells were stained with CD31 and CD45 to identify target cell types: endothelial cells (CD31^+^/CD45^-^) and leukocytes (CD45^+^/CD31^-^). Double negative cells were not further subtyped. Flow cytometry showed an increase in the total leukocyte population in the ipsilateral ischemic brain (**Supplemental Figure 1**), consistent with published observations in the tMCAO mouse model illustrating inflammation in AIS ^27,28^. VCAM-targeted liposomes were preferentially taken up by endothelial cells (**Figure 3A-D**) and leukocytes (**Figure 3E-H**) in tMCAO brains. In the ipsilateral hemisphere, liposome uptake was enhanced in both endothelial cells and leukocytes, compared to the contralateral hemisphere. The number of recovered endothelial cells (**Figure 3C**) and leukocytes (**Figure 3G**) that were liposome-positive trended towards higher values in the ipsilateral hemisphere. Liposome fluorescence intensity in both endothelial cells (**Figure 3D**) and leukocytes (**Figure 3H**) was significantly higher in the ipsilateral hemisphere, as compared to the contralateral.

**Figure 3.**
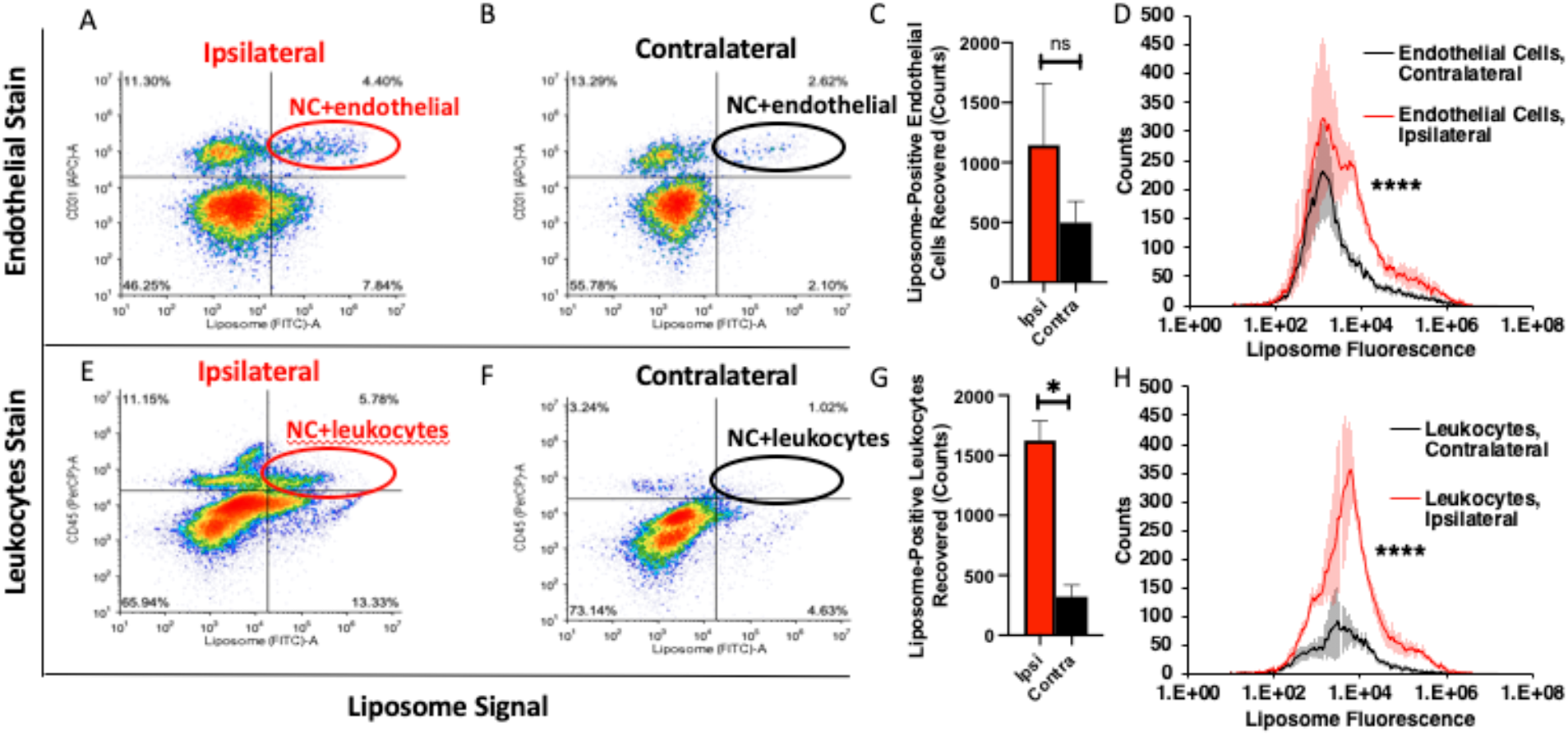
VCAM-targeted liposomes localize to endothelial cells and leukocytes in tMCAO brains. tMCAO mice were IV-injected with fluorophore-labeled VCAM-targeted liposomes (nanocarrier: NC), and single-cell suspensions were prepared from the brains 30 minutes later. Flow cytometry of the ipsilateral hemisphere shows higher VCAM-targeted liposome uptake in endothelial cells (CD31^+^/CD45^-^, A: Ipsilateral, B: Contralateral) and leukocytes (CD45^+^, E: Ipsilateral, F: Contralateral), compared to uptake in the contralateral hemisphere. The contralateral hemisphere had fewer total leukocytes infiltrate into the brain, compared to the ipsilateral hemisphere (E-F, H). Comparison of the contralateral (black) and ipsilateral (red) hemispheres in terms of the number recovered endothelial cells (C) and leukocytes (G) that were liposome-positive showed a trend towards more liposome-positive cells in the ipsilateral hemisphere, for both cell types, with *p<0.05 using student’s t-test. Comparison of liposome fluorescence intensity in endothelial cells (D) and leukocytes (H) of the ipsilateral (red) vs. contralateral (black) brain showed higher liposome uptake in both cell types in the ipsilateral brain, with ***p<0.0001 using Kolmogorov-Smirnov test. N=4.

### VCAM-targeted immunoliposomes deliver small molecule drug to the ischemic brain and reduce infarct volume

Having shown VCAM-targeted NC delivery to brain endothelial cells and leukocytes, we reasoned that VCAM targeting would be well-suited to delivery of therapeutic cargoes for treatment of AIS, especially anti-inflammatory agents ^29^. As our first test drug, we chose dexamethasone-21-phosphate (Dex), a glucocorticoid with anti-inflammatory effects on endothelial cells and leukocytes ^30,31^. Dex is well-established as a treatment for reducing inflammatory brain injury, but was not clinically effective in reducing infarct volume in AIS patients ^32,33^, possibly due to the limited amount of drug delivered to the lesion ^34,35^. By loading Dex into VCAM-targeted liposomes, we hypothesized that it would be concentrated in endothelial cells and leukocytes, ameliorating local inflammation and reducing infarct size in the tMCAO model.

In tMCAO mice, we IV-injected anti-VCAM liposomes loaded with Dex (**Supplemental Table 4**), empty anti-VCAM liposomes, or free Dex at a drug dose of 0.5 mg/kg directly after reperfusion, and then two additional doses at 24-hour intervals. Drug effects were determined at day 3 after reperfusion (**Figure 4A**). In mice that only received IV-injection of the vehicle control, we observed that infarct volume increased by 61% between 24 hours and 72 hours in vehicle controls (**Figure 4B**), as assessed by TTC staining showing metabolically active and inactive tissues, and consistent with previous observations showing the penumbra around the ischemic core slowly dies. No significant effects on infarct size were detected for free Dex or for empty VCAM liposomes, but anti-VCAM liposomes loaded with Dex reduced infarct volume by 35.0 ± 4.8% (**Figure 4C**). Mice receiving free Dex had 50% survival vs. 85% in the Dex loaded VCAM-targeted liposomes (**Figure 4D**). Infarct volume in mice that received anti-VCAM Dex-liposomes at 72 hours was essentially identical to infarct volume at 24 hours post-injury in mice that received the vehicle control (79.8 mm^3^ vs 75.8 mm^3^, *p* = 0.9578). These results suggest that anti-VCAM Dex-liposomes fully arrest infarct volume expansion from 24 to 72 hours.

**Figure 4.**
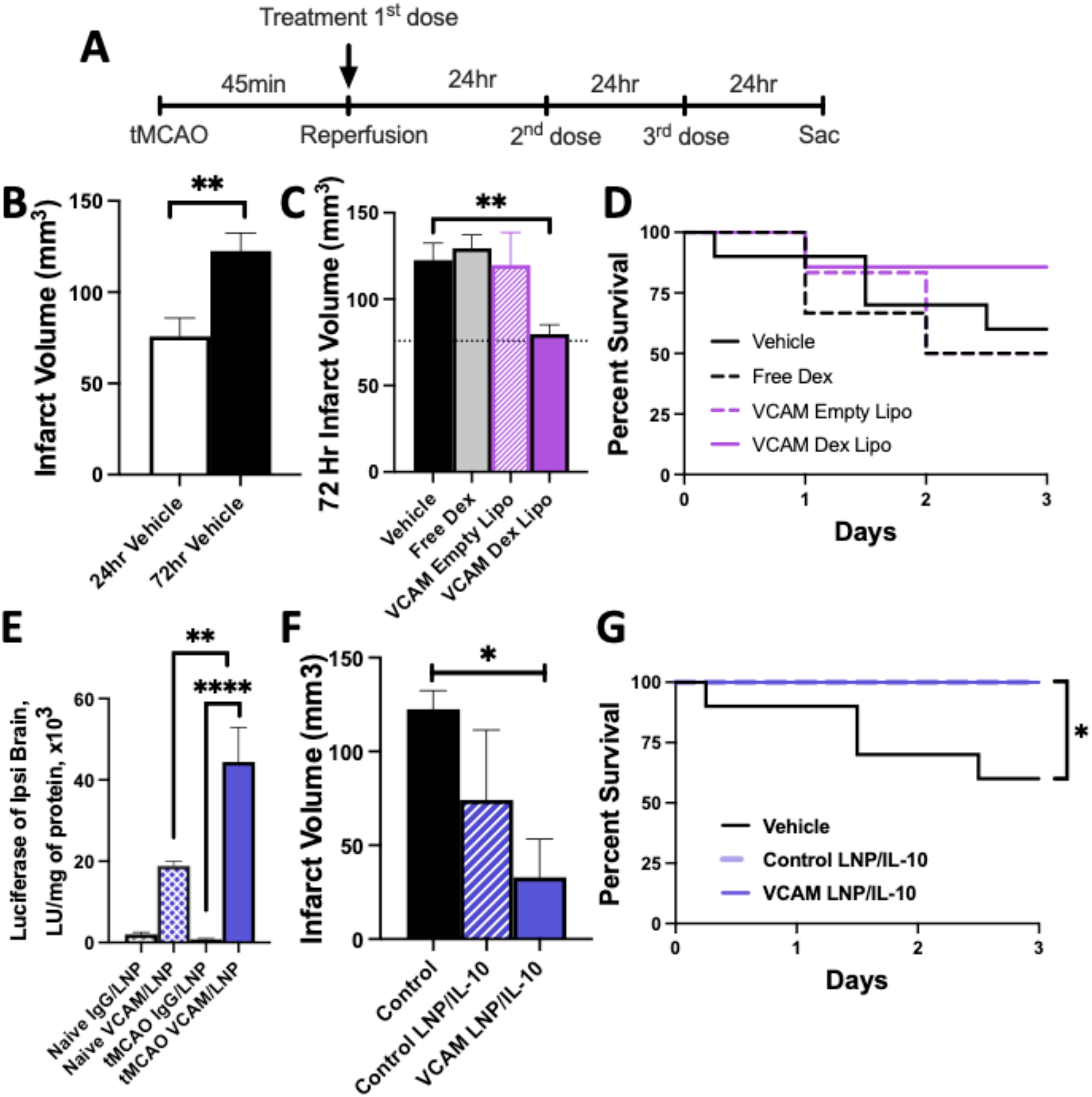
VCAM-targeted NCs deliver diverse cargoes to the brain, and decrease tMCAO-induced infarct size. (A) Mice were treated with VCAM-targeted NCs loaded with the anti-inflammatory drug dexamethasone (Dex) or IL-10 mRNA, or various controls. Each animal received 3 IV injections: one immediately after tMCAO reperfusion, then 2 doses every 24 hours thereafter. Mice were euthanized at 72 hours and the infarct volume was measured post-mortem by staining for dead brain tissue. (B) Stroke volume in tMCAO 24- and 72-hours post-injury. In untreated mice, the stroke volume increased 61.7% between 24 and 72 hours after reperfusion. N=9. **p<0.01 using student t-test. (C) Among the survived animals, VCAM-targeted liposomes loaded with Dex reduced stroke volume at 72 hours, relative to all controls (N=6-10). Neither free Dex (no NC) nor empty VCAM-targeted liposomes decreased stroke volume. (D) Dex-loaded VCAM targeted liposomes showed a higher survival rate (N=6-10). (E) VCAM-targeted LNP delivered mRNA encoding luciferase, producing significantly higher luciferase activity in the ipsilateral hemisphere of tMCAO brains. N=4-9. *p<0.05, **p<0.01, ***p<0.001 using one-way ANOVA followed by Dunnett’s post hoc. (F) IL-10 mRNA-loaded VCAM targeted LNP reduced stroke volume by 73.2%, while untargeted control (bare) LNP had no significant treatment effect. N=4-10. *p<0.05 using one-way ANOVA followed by Dunnett’s post hoc. (G) IL-10 mRNA-loaded VCAM targeted LNP significantly improved survival rate. *p<0.05 using Logrank test.

### VCAM-targeted LNP deliver mRNA to the ischemic brain and ameliorate tMCAO-induced brain injury

LNP have showed great potential in delivering RNA, but little studies have shown therapeutic efficacy of LNP for ischemic stroke. Different from delivery of small molecule drug, which allows potential drug diffusing to cells or surrounding tissue within minutes, the delivery of mRNA provides protection hours after injection that potentially persists for up to 24 days ^36^. We hypothesized that mRNA-loaded VCAM-targeted LNP would allow selective expression in tMCAO brain and outperform untargeted LNP in treatment. We tested functional activity of VCAM-targeted LNP in tMCAO mice after systemic delivery (IV). To assess reporter transgene expression using VCAM-targeted LNP (**Supplemental Table 4**), we IV-injected VCAM-targeted LNP and IgG control LNP containing luciferase mRNA (8µg of mRNA) into tMCAO mice 24 hours after injury and into naïve mice. Four hours after LNP injection, VCAM-targeted LNP produced significantly higher levels of luciferase than untargeted IgG LNP. VCAM targeting improved luciferase expression especially in tMCAO brains (**Figure 4E, Supplemental Figure 2**). Further, we performed studies on cellular transgene expression by VCAM-targeted LNP. LNP were loaded with Cre recombinase-encoding mRNA (**Supplemental Table 4**) to alter fluorophore activity in double-fluorescent mTmG reporter mice ^37^. These mice have a constitutively active Cre-LoxP-flanked tdTomato reporter, followed by a constitutively silent GFP reporter. Expression of Cre recombinase will induce GFP expression in these mice. After 3 LNP doses (8µg mRNA per dose) over 72 hours, VCAM-targeted LNP induced cellular expression of GFP in the ischemic brain (**Supplemental Figure 3**). To our best knowledge, this is the first report describing the delivery of mRNA-LNP in a stroke animal model showing its efficacy in delivering mRNA encoding reporter (luciferase) and/or gene editing proteins (Cre-recombinase).

Encouraged by selective expression of VCAM-targeted LNP in tMCAO brain, we examined the therapeutic effect of these LNP loaded with IL-10 mRNA (**Supplemental Table 4**). IL-10 is a potent anti-inflammatory cytokine and has been reported to down-regulate inflammatory signals and mediate neuroprotection ^38,39^. IV injection of 3 doses of IL-10 mRNA loaded VCAM-targeted LNP (8µg mRNA per dose) after reperfusion (**Figure 4A**) reduced infarct volume by 73.1 ± 16.7%, while the untargeted control (bare) counterpart did not show significant therapeutic effect (**Figure 4F**). Mice receiving IL-10 mRNA LNP had 100% survival (**Figure 4G**). Measurement of IL-10 showed significant elevation of plasma level after IL-10 mRNA loaded LNP (**Supplemental Figure 4**).

## Discussion

Over 1000 drugs have been tested in animal models of AIS and only 1/3 of them showed protective effects preclinically, resulting in over 100 being tested in AIS patients. However, all clinical trials failed to improve outcomes ^12^. The most frequently cited reason for failure was the inability to achieve efficacious concentrations of drugs in brain tissue without causing intolerable side effects. Only recanalizing therapies (thrombolysis and thrombectomies) are efficacious for the treatment of AIS and tPA is the only FDA-approved drug to restore blood flow AIS ^3-5^. Accordingly, the death rate from stroke in the US fell 77% between 1969 and 2013 ^40^. Despite this improvement, 50-55% of stroke patients still become functionally dependent. This raises a central question: Why isn’t it enough to restore blood flow? Sustained inflammation for days to weeks after the initial ischemic event is likely at least part of the answer to this question ^41^. Recent study has shown cerebral microvasculature inflammation upon AIS in both mice and human^42^. We hypothesized that treating sustained inflammation post-AIS may improve stroke outcomes, but to do so, the therapeutic must be efficiently concentrated in the brain.

We aimed to develop nanocarriers (NCs) with the ability to deliver various therapeutics to specific locations in the brain affected by ischemic stroke. Our focus was on utilizing CAMs for this purpose. Previous studies employing microscopy and omics techniques have identified three main types of CAM expression in the brain: A) Pan-endothelial PECAM is constitutively expressed at relatively modest levels; B) ICAM is normally present but upregulated in areas of inflammation and other pathologies; and C) VCAM is modestly expressed under normal conditions but strongly induced in pathological conditions ^21-23^. However, the existing detection methods did not provide detailed information about the accessibility and anchoring capabilities of CAMs for drug delivery systems. To address this, ligands labeled with fluorescence, MRI or isotope probes offered a real-time assessment of the extent, kinetics, and duration of targeting these molecules ^17,25^. Our lab has previously demonstrated the targeting and therapeutic potential of CAM-targeted proteins in animals with brain pathologies, leading to significant anti-thrombotic and anti-inflammatory effects ^43,44^. These results were consistent with the notion that CAMs may provide a promising platform for brain drug delivery.

It is essential to note that previous studies have not thoroughly investigated the accessibility of CAMs for targeted NCs using the chosen model therapeutics. Moreover, the increase in intracranial pressure caused by vasogenic edema can reduce blood flow to the injured brain region, potentially impacting the delivery and local action of the cargo ^45^. Each form of pathological process may have unique effects on drug delivery, and the cargo itself can affect the stability and other features of the drug delivery system. Therefore, our study aims to address these challenges and provide novel insights into drug delivery system design, considering the role of the pathology and other factors involved.

Here, we demonstrated delivery to the ischemic hemisphere of IV-injected mAbs and the two most clinically relevant targeted NCs (liposomes and LNP) directed to different CAMs. Targeting *to* the BBB via endothelial surface marker CAMs allows us to develop intervention strategies that correct BBB deficits by restoring its function and preventing further damage against toxic plasma proteins^46^. We tested targeting to PECAM, ICAM and VCAM. Comparing targeted vs. non-targeted delivery to the AIS-injured region of the brain, CAM targeting yielded 37-120-fold increased uptake for mAbs, 4-16-fold increased uptake for LNP, and 11-50-fold increased uptake for liposomes. In all cases, VCAM targeting yielded the highest uptake in the injured hemisphere of the brain. This positions VCAM as the most attractive target for drug delivery in AIS.

Agreeing with previous studies, flow cytometry analysis of brains from tMCAO mice showed a marked influx of leukocytes in the injured brains ^47^. VCAM-targeted liposomes bound to endothelial cells, in agreement with prior studies ^17^; however, they also accumulated in leukocytes in this model. This delivery feature is advantageous, as overzealous leukocytes in the injured region of the brain after stroke are known to be associated with poor patient outcomes ^48,49^. By providing delivery to the two main cell types driving post-stroke inflammation – endothelial cells and leukocytes, VCAM-targeted NCs are likely able to address this underlying mechanism of secondary injury and protracted cell death that occurs in the penumbra around the ischemic core ^50^. Knowledge of the cell types responsible for delivery to the brain in tMCAO mice was useful in making informed decisions for what drug to test as a therapeutic.

We demonstrated that VCAM-targeted NCs can be used to deliver pharmacologically-relevant concentrations of small molecule drugs (liposomes) and mRNA (LNP), showing the versatility of this targeting strategy (**Figure 4**). Small molecule drugs can be delivered and act on specific cells within minutes, as well as diffuse to surrounding cell and tissue to provide protection. Thus, it is considered as “early protection”. On the other hand, transcription and protein synthesis after mRNA delivery takes hours but the effect can last from 24 hours to days, which is considered as “delayed protection”. To attack post-stroke inflammatory injury, we selected anti-inflammatory drugs Dex and mRNA encoding for IL-10 as a cargo drugs to be used with VCAM-targeted NCs. Dex is a glucocorticoid with anti-inflammatory effects on both endothelial cells and leukocytes, and is relatively easy to encapsulate into targeted NCs [12, 30]. It has effects on both endothelial cells and leukocytes by downregulating inflammatory signals ^51^, potentially transforming the pro-inflammatory environment of the ischemic penumbra into an anti-inflammatory one. Dex was previously tested as a free drug in clinical trials to treat AIS but failed due to lack of clear therapeutic benefits and it induced severe systemic side effects ^52,53^. For all these reasons, we loaded Dex into VCAM-targeted liposomes. We found that VCAM-targeted Dex-liposomes reduced infarct volume by 35% vs. vehicle control and free drug (*p* = 0.02 and 0.049, respective). In contrast to the free drug, we observed no brain hemorrhagic transformation and no gastrointestinal bleeds (**Supplemental Figure 5**).

Interleukin 10 (IL-10) is a potent cytokine with multifaceted anti-inflammatory properties that play a crucial role in the resolution of inflammation after AIS. IL-10 has been investigated in clinical trials for its potential therapeutic use in chronic inflammatory diseases. In the context of AIS, IL-10 is essential for resolving the inflammatory phase and promoting neuronal and glial cell survival ^38,54^. Human studies have indicated that low plasma levels of IL-10 are associated with poorer prognoses and outcomes in patients with AIS ^55^. Our research findings demonstrate that using VCAM-targeted LNP encapsulating IL-10-encoding mRNA resulted in a 73% reduction in infarct volume. Additionally, we observed a 100% survival rate with no bleeding or lung damage in mice receiving this treatment. Untargeted (control) LNP carrying IL-10 mRNA also resulted in 100% survival, but with a bimodal distribution in infarct volume, indicating responders and non-responders. We also observed that 25% of mice treated with untargeted LNP experienced lung injury and aqueous intestines, in contrast to the targeted LNP group bleeds (**Supplemental Figure 6**).

Among drugs that made it to AIS clinical trials, the average improvement in infarct volume in rodents was 25% ^12^, so our result with VCAM targeting is promising. In fact, VCAM-targeted Dex-liposomes brought the infarct volume at 72 hours down to the same level observed at 24 hours in mice that received vehicle only. This suggests that VCAM-targeted Dex-liposomes can abrogate neuronal death in the inflammatory penumbra. In addition, we observed that we can target LNP to the ischemic brain for RNA delivery, which opens a new, nearly endless set of opportunities for delivery of either mRNA encoding for therapeutic proteins without the need to significantly optimize LNP formulations.

In summary, this study uses the tMCAO model of AIS to show that targeting CAMs, especially VCAM, with mAbs and NCs is a promising direction for improvement of AIS therapies. VCAM-targeted NCs and mAbs accumulate in the AIS brain at levels 1-2 orders of magnitude higher than untargeted controls. VCAM targeting delivers NCs to key cell types involved in post-stroke inflammation, namely endothelial cells and leukocytes. VCAM-targeted LNP are able to transfect and express mRNA in the ischemic brain. VCAM-targeted Dex-loaded liposomes reduced stroke size to a greater extent than most drugs that have moved from rodent to human studies. Thus, VCAM-targeted therapeutics represent an attractive platform for treating AIS.

## Acknowledgement

J.N. received funding from the American Heart Association (Grant 916172). O.A.M.-C. received funding from the American Heart Association (Grant 19CDA34590001). P.M.G. received funding from the National Institutes of Health K99 (K99HL153696). M.E.Z. received funding from the National Institutes of Health F31 (F31 HL154662-01). S·O. received funding from the American Heart Association (Grant 23PRE1014444). V.R.M and J.S.B. received support from the Cardiovascular Institute of the University of Pennsylvania. J.S.B. received funding from K08-HL-138269, R01-HL-153510, R01-HL-160694, R01-HL-157189, R21-AI-166778-01. V.R.M received funding from the National Institutes of Health (NIH) (R01 HL155106, R01 HL128398, R01 HL143806).

## Disclosures

Declaration of Competing Interest J.S.B., V.R.M., and O.A.M-C. have pending patent applications fully disclosed by the University of Pennsylvania. Other authors report no financial disclosures.

### Supplemental Material

Figure S1-S6

Table S1-S5

