## Supplemental Figures and Tables for "Targeting lipid nanoparticles to the blood brain barrier to ameliorate acute ischemic stroke"

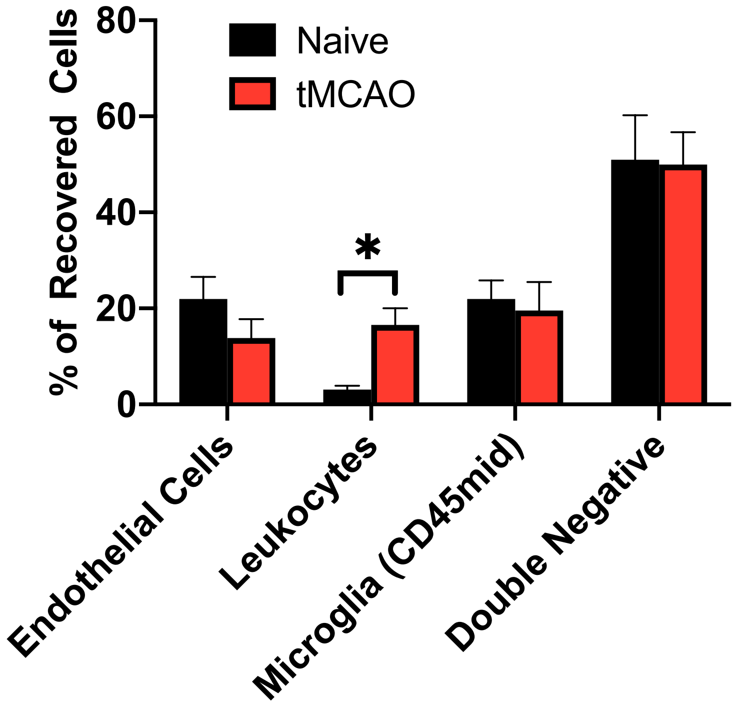


**Supplemental Figure 1.** Flow cytometry analysis of cell recovery in ipsilateral hemisphere of naïve (corresponding side) vs tMCAO brain showed significant infiltration of leukocytes after injury. Data reported as mean ± SEM. N = 3/group.

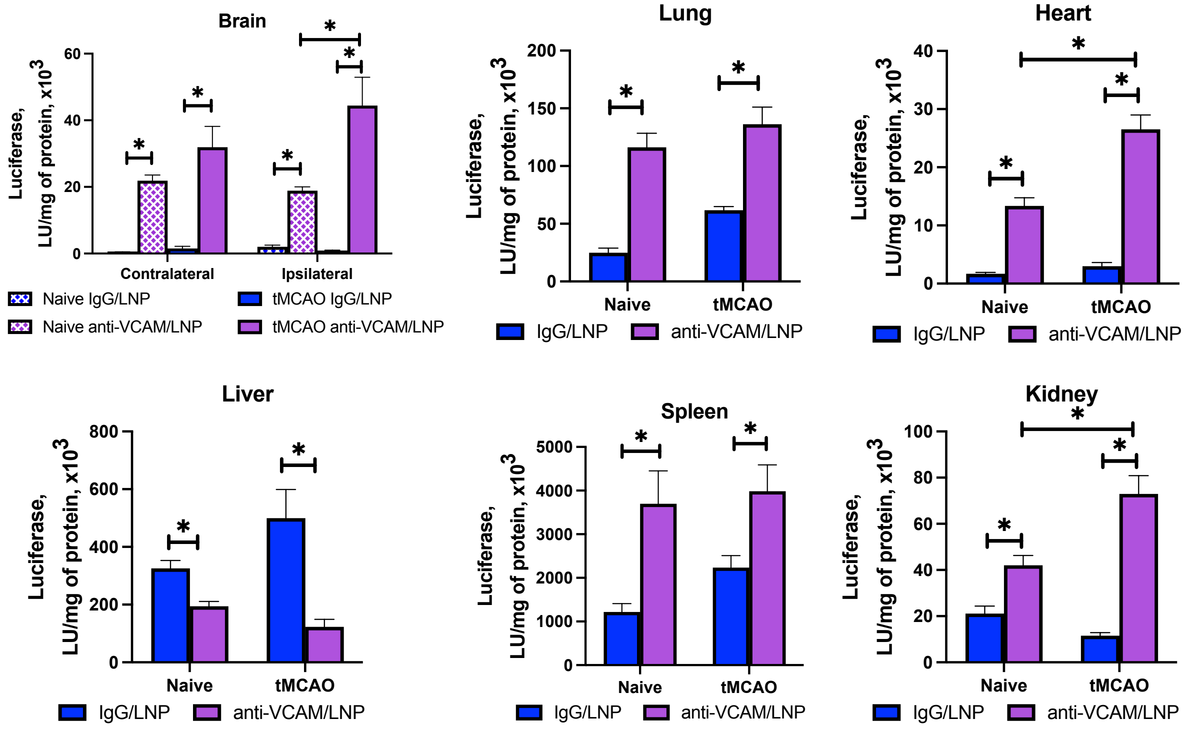


**Supplemental Figure 2.** Luciferase expression in naïve vs tMCAO brain and other major organs. Data reported as mean ± SEM. N = 4/group. P<0.05 using one-way ANOVA with Dunnett's post-hoc or student's t-test.

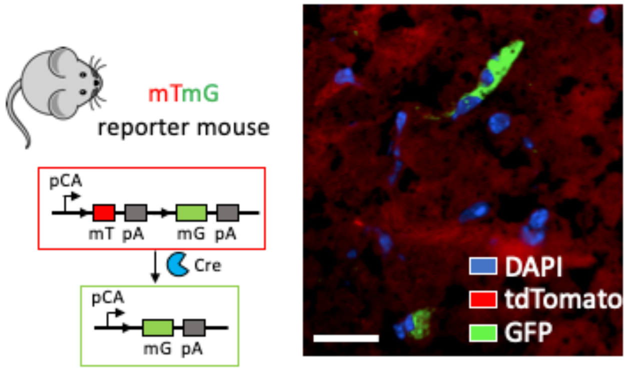


**Supplemental Figure 3**. Histology of brain slice from a mTmG mouse injected with LNP-Cre mRNA. GFP signal indicates mG-positive cells that were successfully transfected by the LNP. Scale bar: 25 µm.

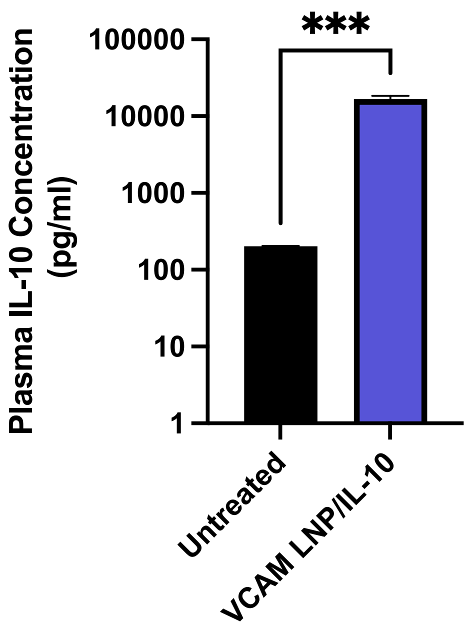


**Supplemental Figure 4**. Treatment of VCAM-LNP loaded with IL-10 mRNA significantly elevated plasma IL-10 concentration. Data reported as mean ± SEM. N = 3-4/group.

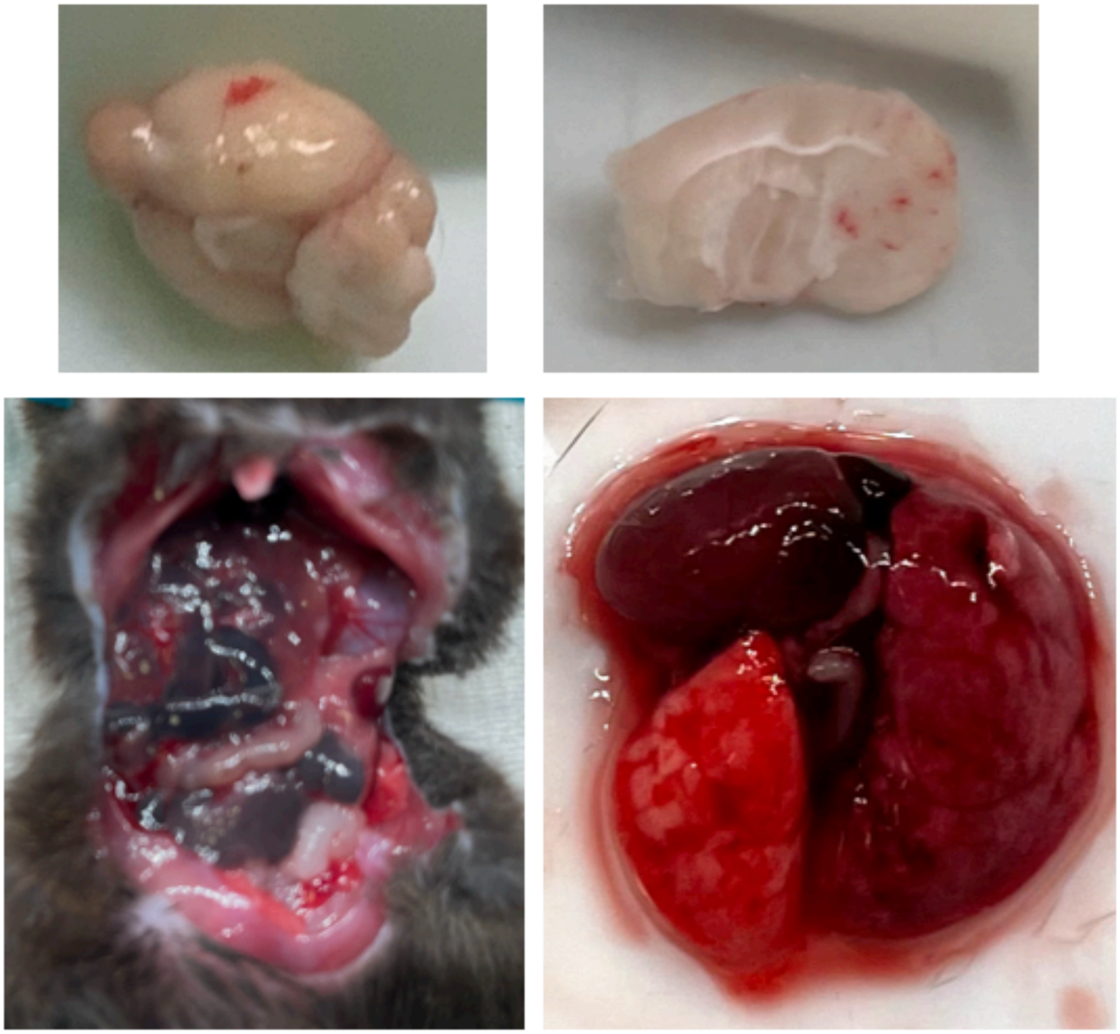

**Supplemental Figure 5**. Hemorrhagic transformation, gastrointestinal bleeding and lung injury after free dexamethasone treatment.

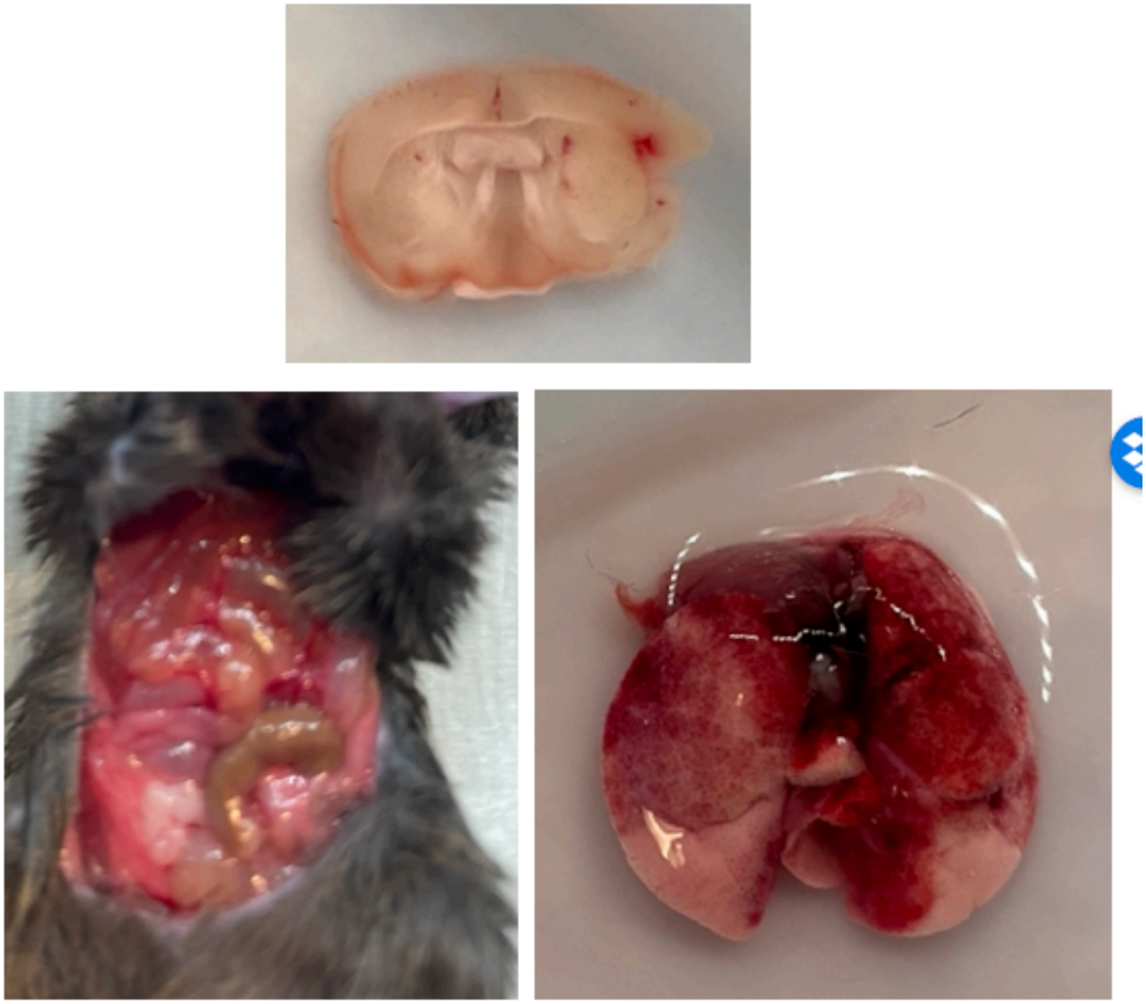

**Supplemental Figure 6**. Hemorrhagic transformation, aqueous intestines and lung injury after control IL-10 LNP treatment.

**Supplemental Table 1.** Biodistribution of anti-CAM mAbs injected 24 hours after tMCAO. Experimental design as in **Figure 1B**. Values reported as percent of injected dose/g organ (%ID/g). Samples were collected 30 minutes after IV injection. Data reported as mean ± SEM. N = 3/group. *p<0.05 compared to IgG using one-way ANOVA with Dunnett post-hoc. (Ipsi: ipsilateral hemisphere of tMCAO brain; Contra: contralateral hemisphere)

| mAb | Blood | Lung | Liver | Spleen | Contra | Ipsi |
|---|---|---|---|---|---|---|
| **IgG** | 41.0±0.8 | 5.2±3.5 | 9.8±1.7 | 9.7±3.9 | 0.1±0.05 | 0.3±0.03 |
| **αPECAM** | 8.5±0.8* | 73.4±2.9* | 27.5±0.9* | 19.9±2.5 | 2.34±0.58* | 2±0.82 |
| **αICAM** | 6.7±0.9* | 197.5±12.5* | 21.6±0.6* | 24.9±2.2 | 0.82±0.12 | 1.38±0.21 |
| **αVCAM** | 5.4±0.3* | 10.9±1.1 | 22.8±2.3* | 106.2±13.5* | 2.45±0.33* | 3.95±0.87* |

**Supplemental Table 2.** Biodistribution of anti-CAM LNPs injected 24 hours after tMCAO. Experimental design as in **Figure 1B**. Values reported as percent of injected dose/g organ (%ID/g). Samples were collected 30 minutes after IV injection. Data reported as mean ± SEM.

N = 4-6/group. *p<0.05 compared to IgG using one-way ANOVA with Dunnett post-hoc. (Ipsi: ipsilateral hemisphere of tMCAO brain; Contra: contralateral hemisphere)

| LNPs | Blood | Lung | Liver | Spleen | Contra | Ipsi |
|---|---|---|---|---|---|---|
| **IgG** | 10.1±1.3 | 9.5±0.6 | 43±3.8 | 27.3±4.3 | 0.24±0.03 | 0.25±0.01 |
| **αPECAM** | 7.4±0.3 | 85±2.2* | 29.8±0.2* | 50.5±3.1 | 1.07±0.07* | 1.30±0.04* |
| **αICAM** | 8.8±0.8 | 172.7±21* | 24.9±1.8* | 48.5±7.2 | 0.5±0.09 | 0.89±0.09 |
| **αVCAM** | 7.8±0.9 | 30.7±3.3 | 24.4±1.9* | 113.8±7* | 1.84±0.17* | 2.83±0.26* |

**Supplemental Table 3.** Biodistribution of anti-CAM liposomes injected 24 hours after tMCAO. Experimental design as in **Figure 1B**. Values reported as percent of injected dose/g organ (%ID/g). Samples were collected 30 minutes after IV injection. Data reported as mean ± SEM. N = 4-6/group. *p<0.05 compared to IgG using one-way ANOVA with Dunnett post-hoc. (Ipsi: ipsilateral hemisphere of tMCAO brain; Contra: contralateral hemisphere)

| Liposomes | Blood | Lung | Liver | Spleen | Contra | Ipsi |
|---|---|---|---|---|---|---|
| **IgG** | 19.1±1.7 | 12.8±0.8 | 40±2.3 | 32.1±1.9 | 0.2±0.02 | 0.3±0.04 |
| **αPECAM** | 5.8±2.2* | 99.1±3* | 30.8±4.4 | 43.2±1.2 | 0.6±0.1* | 1.2±0.1* |
| **αICAM** | 3.7±0.2* | 171.2±18.8* | 24.4±0.8 | 36.2±6.1 | 1.3±0.06* | 1.0±0.12* |
| **αVCAM** | 2.1±0.3* | 14.3±2.4 | 43.9±5.9 | 78.9±13.1* | 1.0±0.05* | 1.9±0.2* |

**Supplemental Table 4. Characterization of targeted nanocarriers.** N=3. All values represented as mean ± SEM of three independent measurements.

| | **VCAM-Dex-liposome** | **IgG-luciferase LNP** | **VCAM-luciferase LNP** | **VCAM-Cre recombinase LNP** | **Control IL-10 LNP** | **VCAM-IL-10 LNP** |
|---|---|---|---|---|---|---|
| **Size** | 161 ± 2 | 111 ± 0.1 | 115 ± 0.1 | 107 ± 3 | 78 ± 0.25 | 102.5 ± 0.9 |
| **PDI** | 0.15 ± 0.032 | 0.09 ± 0.006 | 0.10 ± 0.025 | 0.08 ± 0.012 | 0.043 ± 0.009 | 0.116 ± 0.014 |
| **Drug Loading** | 13.8 ± 0.7% | 97% | 97% | 97% | 97% | 97% |

**Supplemental Table 5. Antibodies used for flow cytometry**

| **Surface Marker** | **Clone** | **Fluorophore** |
|---|---|---|
| CD31 | MEC13.3 | Alexa fluor-647 |
| CD45 | 30-F11 | PerCP/Cyanine5.5 |
